# Evaluation of deep-learning-based lncRNA identification tools

**DOI:** 10.1101/683425

**Authors:** Cheng Yang, Man Zhou, Haoling Xie, Huaiqiu Zhu

## Abstract

Long non-coding RNAs (lncRNAs, length above 200 nt) exert crucial biological roles and have been implicated in cancers^1,2^. To characterize newly discovered transcripts, one major issue is to distinguish lncRNAs from mRNAs. Since experimental methods are time-consuming and costly, computational methods are preferred for large-scale lncRNA identification. In a recent study, Amin et al.^3^ evaluated three deep-learning-based lncRNA identification tools (i.e., lncRNAnet^4^, LncADeep^5^, and lncFinder^6^) and concluded “The LncADeep PR (precision recall) curve is just above the no-skill model and LncADeep showed poor overall performance”. This surprising conclusion is based on the authors’ use of a non-default setting of LncADeep. Actually, LncADeep has two models, one for full-length transcripts, and the other for transcripts including partial-length. Being aware of the difficulty of assembling full-length transcripts from RNA-seq dataset, LncADeep’s default model is for transcripts including partial-length. However, according to the results posted on Amin et al.’s website, the authors used LncADeep with full-length model, while they claimed to use the default setting of LncADeep, to identify lncRNAs from GENCODE dataset, which is composed of full- and partial-length transcripts. Thus, in their evaluation, the performance of LncADeep was underestimated. In this correspondence, we have tested LncADeep’s default setting (i.e., model for transcripts including partial-length) on the datasets used in Amin et al.^3^, and LncADeep achieved overall the best performance compared with the other tools’ results reported by Amin et al.

---

To compare our new evaluation of LncADeep with Amin et al.’s reported results, we follow their evaluation metrics, among which “accuracy” and “F1 score” captures the overall performance of a tool. Table 1 displays the performance of lncRNA identification in both human and mouse datasets. In human dataset (denoted by lncGH), LncADeep achieves superior performance, with the highest accuracy of 94.6% (2.3% higher than that of lncRNAnet and 10.4% higher that of lncFinder), F1 score of 89.2% (4% higher than that of lncRNAnet and 16.1% higher than that of lncFinder), precision of 89.2%, recall of 97.5% and the lowest error rate of 5.4%, while lncRNAnet only shows a higher specificity. In mouse dataset (denoted by lncGM), compared with other tools, LncADeep still achieves the best performance in all metrics except specificity. Although lncFinder’s specificity is the highest, its precision and recall are much lower than LncADeep’s, and leads to the accuracy (89.5%) and F1 score (79.8%) of lncFinder evidently lower than those of LncADeep (accuracy of 95% and F1 score of 89.3%). Apart from the above metrics, we plot ROC and PR (precision recall) curves of lncRNA identification, where LncADeep has the highest AUC (Fig. 1) and AP (Fig. 2), demonstrating that LncADeep yields overall the best performance for lncRNA identification on human and mouse datasets compared with lncRNAnet and lncFinder.

**Table 1.** The lncRNA identification performance of lncRNAnet, LncADeep and lncFinder on human and mouse datasets.

| Metrics<sup>a</sup> | lncGH<sup>b</sup> |  |  | lncGM<sup>b</sup> |  |  |
| --- | --- | --- | --- | --- | --- | --- |
|  | lncRNAAnet<sup>c</sup> | LncADeep<sup>d</sup> | lncFinder<sup>c</sup> | lncRNAAnet<sup>c</sup> | LncADeep<sup>d</sup> | lncFinder<sup>c</sup> |
| Accuracy | 92.3% | **94.6%** | 84.2% | 77.3% | **95.0%** | 89.5% |
| Precision | 75.8% | **82.1%** | 59.3% | 48.1% | **83.2%** | 68.6% |
| Recall | 97.2% | **97.5%** | 95.2% | 62.5% | **96.4%** | 95.2% |
| F1 score | 85.2% | **89.2%** | 73.1% | 54.4% | **89.3%** | 79.8% |
| Specificity | **90.9%** | 82.1% | 80.9% | 81.4% | 83.2% | **87.9%** |
| Error rate | 7.6% | **5.4%** | 22.5% | 22.6% | **5.0%** | 10.4% |

(a) Evaluation metrics are defined as follows: Accuracy=(TP+TN)/(TP+TN+FP+FN),
Precision=TP/(TP+FP), Recall=TP/(TP+FN), Specificity=TN/(TN+FP),
F1 score=2×Precision×Recall/(Precision+Recall),
Error rate=(FP+FN)/(TP+TN+FP+FN), where TP, TN, FP and FN represents true positive, true negative, false positive and false negative, respectively. lncRNA is treated as positive. Recall measures the ratio of actual positives which are correctly identified, precision measures the ratio of true positives in all predicted positives, specificity measures the ratio of actual negatives which are correctly identified, F1 score and accuracy are composite measures used as aggregated performance scores for the evaluation of algorithms.
(b) lncGH and lncGM refers to GENCODE human and mouse dataset respectively, which have been used to evaluate the performance of lncRNA identification tools by Amin et al. The ratio of lncRNA to mRNA in lncGH (lncGM) is around 1:3, which is class-imbalance.
(c) The results of lncRNAAnet and lncFinder are from Amin et al.
(d) LncADeep's default model (i.e., model for transcripts including partial-length) is used for lncRNA identification.

**Figure 1.**
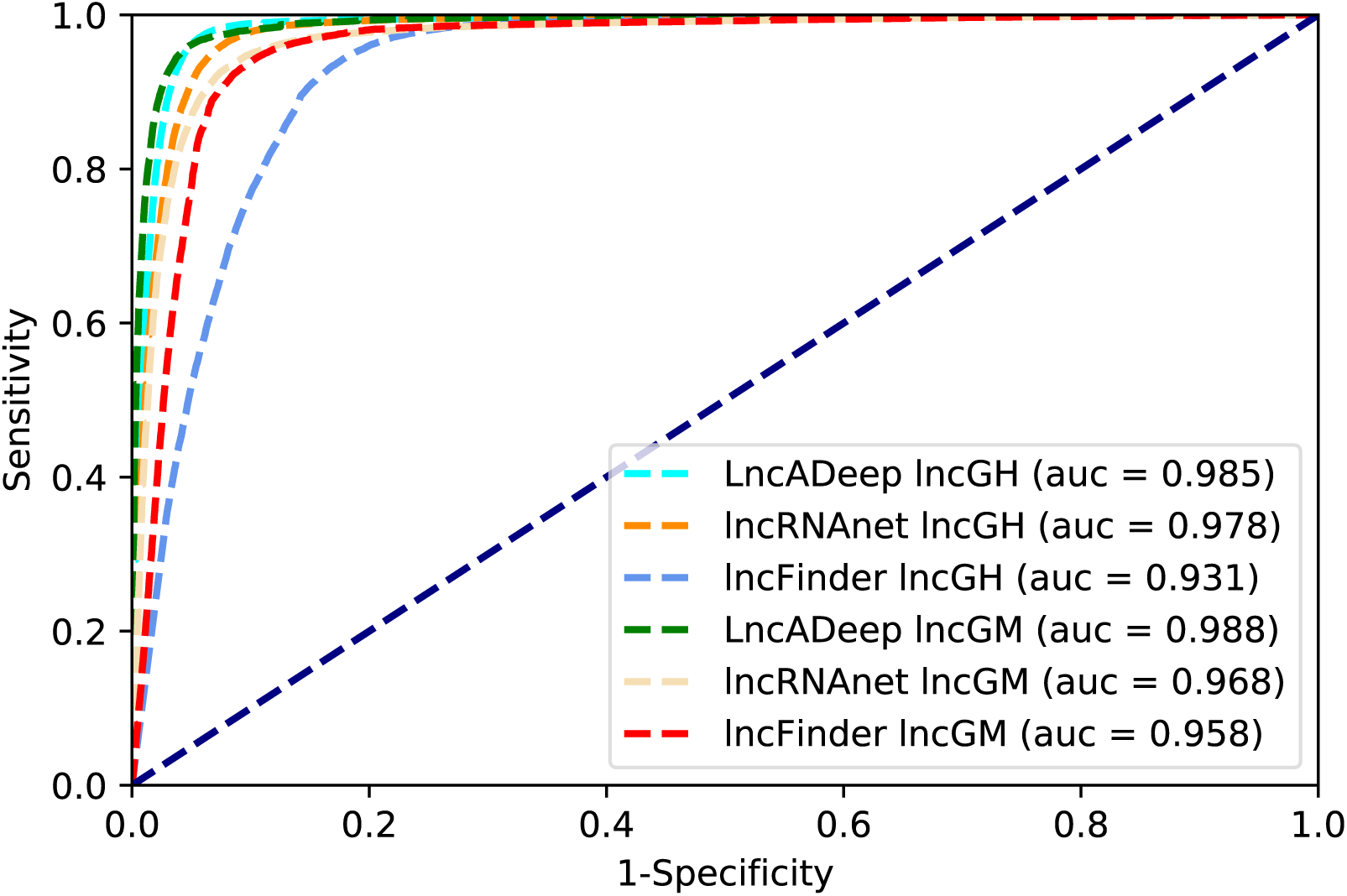
lncRNA identification ROC curves for lncRNAnet, LncADeep and lncFinder. (a) lncGH and lncGM refers to GENCODE human and mouse dataset respectively, which are from Amin et al. ROC curve measures the discrimination capability of a tool, especially when the test dataset is class-balance. The higher area under the curve (auc), the better.

**Figure 2.**
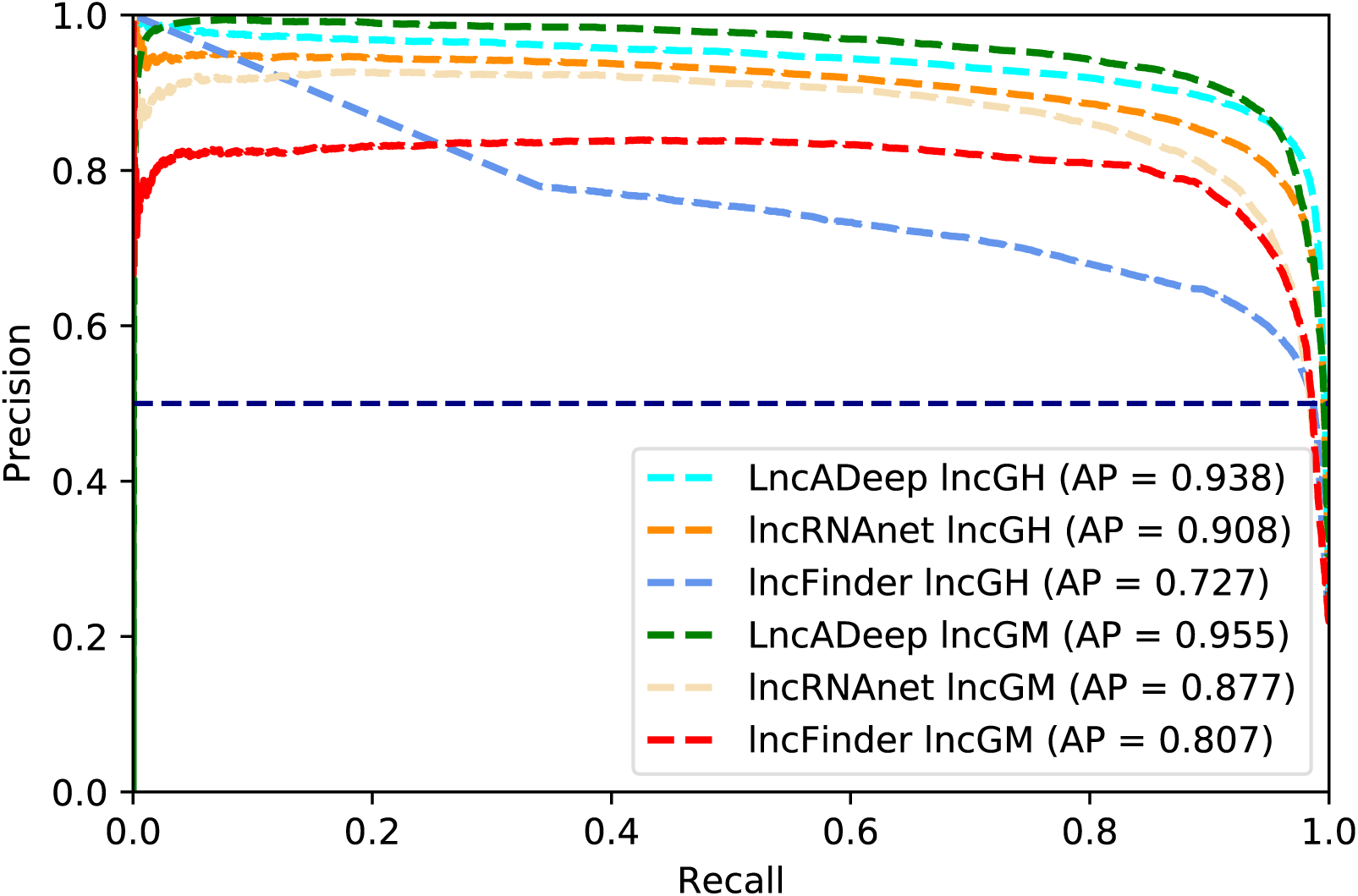
lncRNA identification precision recall (PR) curves for lncRNAnet, LncADeep and lncFinder. (a) lncGH and lncGM refers to GENCODE human and mouse dataset respectively, which are from Amin et al. PR curve measures the discrimination capability of a tool, especially when the test dataset is class-imbalance. The higher area under the PR curve (AP), the better.

As discussed previously in LncADeep^5^, currently, most RNA-seq data are based on the second-generation sequencing technologies, tending to produce short reads which impede the accurate full-length transcript assembly. It has been estimated that over 60% of transcripts reconstructed by the best-performing unguided assembly tools are of partial-length^7^. Partial-length mRNA truncated at 5’ and/or 3’ end can lead to incomplete coding sequence (CDS) and incline to be misclassified as lncRNA, which further complicates lncRNA identification. Therefore, LncADeep proposed the model targeting on transcripts including partial-length and the test results^5^ have demonstrated its effectiveness. With the development of sequencing technologies, third-generation sequencing technologies, such as PacBio and Nanopore, can produce full-length transcripts^8^. For this scenario, LncADeep is suitable for lncRNA identification with its full-length model.

In conclusion, we have carried out new evaluation of LncADeep on lncRNA identification to investigate the surprising claim that “The LncADeep PR curve is just above the no-skill model and LncADeep showed poor overall performance^3^.” Our evaluation shows that LncADeep actually performs quite well for lncRNA identification, while Amin et al. used a non-default setting (i.e., model for full-length transcripts) of LncADeep to identify lncRNAs from transcripts including partial-length and much underestimated LncADeep. Although the evaluation of deep-learning-based lncRNA identification tools by Amin et al. is of importance, it should be noted that the setting of lncRNA identification tools need be appropriate and clearly mentioned, otherwise a state-of-the-arts tool could be underestimated and academic users be misled.

## Data availability

The evaluation results of LncADeep are available at https://github.com/cyang235/lncadeep.results/.

## Author contributions

All authors contributed to the manuscript. HZ and CY conceived the idea. MZ performed the experiment. CY wrote the paper. HX and HZ revised the manuscript.

## Competing interests

The authors declare no competing interests.

## Notes

https://github.com/cyang235/lncadeep.results/

